# PaGeSearch: A Tool for Identifying Genes within Pathways in Unannotated Genomes

**DOI:** 10.1101/2023.09.26.559665

**Authors:** Sohyoung Won, Jaewoong Yu, Heebal Kim

**Affiliations:** Department of Agricultural Biotechnology and Research Institute of Agriculture and Life Sciences, Seoul National University, Seoul, Republic of Korea; UNGENE, Inc, Seoul, Republic of Korea; eGnome, Inc, Seoul, Republic of Korea

## Abstract

In biological research, the identification and comparison of genes within specific pathways across the genomes of various species are invaluable. However, annotating the entire genome is resource-intensive and sequence similarity searches often yield results that are not actually genes. To address these limitations, we introduce PaGeSearch (Pathway Gene Search), a tool designed to identify genes from predefined lists, especially those in specific pathways, within genomes. The tool employs an initial sequence similarity search to identify relevant genomic regions, followed by targeted gene prediction and neural network-based result filtering. PaGeSearch suggests the regions that are most likely the orthologs of the genes in the query and is designed to be applicable for species within five classes: mammals, fish, birds, eudicotyledons, and liliopsida. When compared with GeMoMa, PaGeSearch consistently outperformed in terms of sensitivity and positive predictive value. Although its performance showed increased variability when applied to actual biological pathways, it nonetheless maintained an acceptable level of accuracy.

## INTRODUCTION

Advances in next-generation sequencing (NGS) have led to the production of a vast number of eukaryotic genome sequences. These genome sequences and assemblies can be easily accessed and downloaded from various databases, such as NCBI or Ensembl. However, the extent of post-assembly processing varies across genomes. For some species, chromosome-level assemblies and annotation information are readily available, whereas for others, only contig-level assemblies exist, and annotation remains incomplete.

Genome annotation, the process of identifying genes and other functional elements within a genome sequence, is a crucial step in analyzing and comparing genomes. In order to study the genes present within a species’ genome or to compare gene sequences between species, it is essential to identify and describe the genes.

Several genome annotation tools have been developed to date, such as GeneMark-EP+ (Brůna et al. 2020), MAKER (Cantarel et al. 2008), and BRAKER (Hoff et al. 2019), have significantly enhanced the efficiency and accuracy of genome analysis. Despite these advancements, the substantial amount of data involved, including genome sequences, expressed sequence tags (ESTs), and protein databases, still demands considerable computational resources and time.

In some cases, researchers may be interested in a specific group of genes, such as those in a particular pathway, from one or multiple species to study their conservation, evolution, or functional significance. In these situations, performing whole-genome annotation of various species can be computationally inefficient and time-consuming. To address this problem, sequence alignment (or searching) tools like BLAST (McGinnis and Madden 2004), HMMER (Eddy 1998), and Exonerate (Slater and Birney 2005) are often employed. However, these tools have limitations, as they can only find similar parts within gene sequences and often fail to identify the complete sequence of the gene, moreover, they cannot confirm whether the identified regions are actually part of the gene in question.

To tackle these issues, alternative tools like genBlastA (She et al. 2009) and genBlastG (She et al. 2011) were developed to enhance the identification of homologous genes from BLAST search results. GenBlastA employs a graph-based algorithm to filter high-scoring pairs (HSPs) into well-defined groups, each of which represents a candidate gene in the target genome. It also introduces a novel edge length metric for identifying gene boundaries. GenBlastG takes the groups of HSPs identified by genBlastA and determines gene models by maximizing the similarity to their corresponding queries while examining alignments and neighboring genomic regions for start/stop codons and splicing signals. However, genBlastA and genBlastG rely on BLAST, which can be time-consuming compared to more recently developed and efficient alternatives, such as mmseqs2 (Steinegger and Söding 2017) or DIAMOND (Buchfink et al. 2015). GeMoMa (“Gene Model Mapper”) also offers an option for predicting a list of reference transcripts in newly sequenced genomes (Keilwagen et al. 2019). GeMoMa is a gene prediction tool that leverages knowledge from well-annotated reference genomes and features an option called “selected,” which filters the transcripts in a given list and annotates only relevant sections of the genome.

Biological pathways are networks of interconnected genes and molecules that coordinate and regulate various biological processes within cells, tissues, and organisms. By identifying the genes present in a pathway, researchers can gain insights into their collective functions and how they interact with each other to carry out specific biological processes. This knowledge can help elucidate the underlying molecular mechanisms and regulatory networks that govern cellular processes, providing a deeper understanding of biological systems.

Here, we present PaGeSearch, a tool that is designed to effectively identify a list of genes within a genome, with a focus on genes associated with specific pathways. Utilizing an initial sequence similarity search to identify candidate regions, and subsequently performing gene prediction within these regions, PaGeSearch significantly narrows down the search space. This targeted approach enables more efficient calculations and mitigates computational burden by only focusing on relevant genomic subsets, as opposed to the entire sequence. Following this, PaGeSearch uses a neural network model to provide candidates that are the most likely orthologs of the query genes. Only the assembly and query gene sequences are required to run PaGeSearch. The tool also includes a pipeline for downloading orthologous gene sequences for the pathway of interest from the Ensembl database, which can be used as the searching query. PaGeSearch is designed to be applicable across five taxonomic classes: mammals, fish, birds, eudicotyledons, and liliopsida, and is calibrated based on the genome and gene models of well-annotated model organisms within each class.

We investigated the performance of PaGeSearch on both the genomes of archetype species and those of related species. Furthermore, we compared the tool’s performance to that of other competing software, such as GeMoMa. Finally, we applied PaGeSearch to real-world pathway gene identification problems, demonstrating its particular utility for comparative analyses of genes across different species.

## RESULTS

### Overview of PaGeSearch

The pipeline of PaGeSearch consists of three major parts as illustrated in Figure 1; (1) defining candidate regions, (2) gene prediction at the candidate regions, and (3) validating and filtering the gene prediction results. There are three steps for defining candidate regions; (i) MMseqs2 searching in tblastn mode (Steinegger and Söding 2017), and (iii) grouping and extending the MMseqs2 search results. Gene prediction at the candidate regions is achieved in two steps; (i) making protein hints from the query protein sequences using Exonerate (Slater and Birney 2005), and (ii) ab initio gene prediction using Augustus (Stanke et al. 2006). To validate and filter the gene prediction results, we employed two steps; (i) aligning the query sequences to the gene predicted sequences with Exonerate, and (ii) filtering the results based on scores calculated from protein alignment and gene prediction metrics using neural network models pre-trained for each class; mammals, fish, birds, eudicotyledons, and liliopsida.

**Figure 1.**
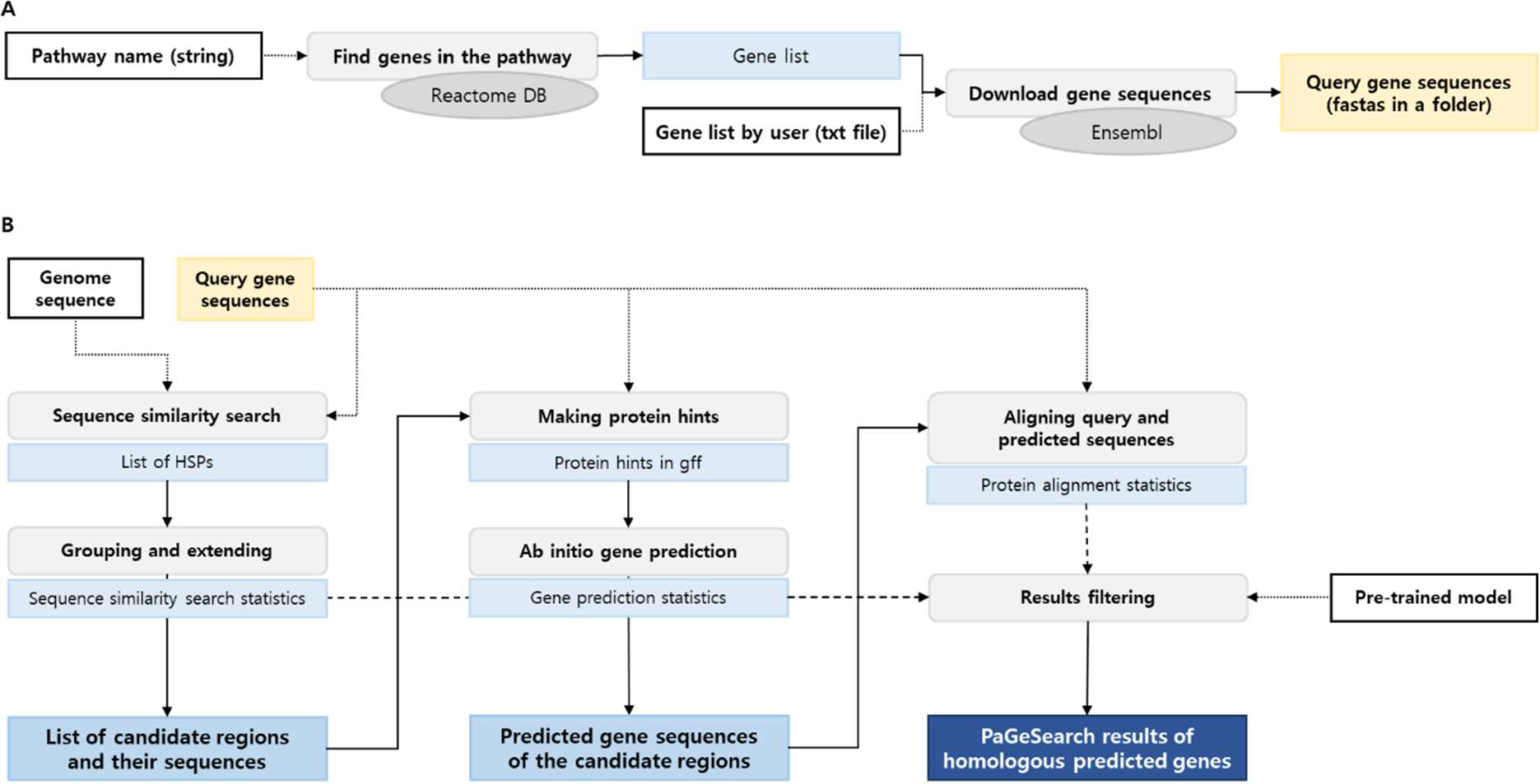
Process of PaGeSearch. (A) Genes in the specified pathway are identified from the Reactome database, and their sequences are downloaded from the Ensembl database for use as the PaGeSearch query. (B) PaGeSearch initiates by finding seed regions through sequence similarity search, followed by gene prediction within these regions. The final results are filtered using a neural network model that evaluates sequence similarity, gene prediction metrics, and protein alignment statistics.

### Benchmark

We evaluated the performances of PaGeSearch and a comparable existing tool, GeMoMa (Figure 2). GeMoMa can predict genes corresponding to a given list of transcripts, from reference genomes and GFFs (Keilwagen et al. 2019). We used the “GeMoMaPipeline” and provided the list of transcript IDs corresponding to the query genes in the “selected” parameter. We used all transcript IDs that match the gene ID from Ensembl as the input. Default values were used for the remaining parameters. We assessed the sensitivity, specificity, and positive predictive value (PPV), and negative predictive value (NPV) of PaGeSearch and GeMoMa in the four species used for model construction (‘Same’, ‘Within Order’, Within Class’, ‘Within Phylum’) and a different test species (‘Test’), respectively (Table 1).

**Table 1.**
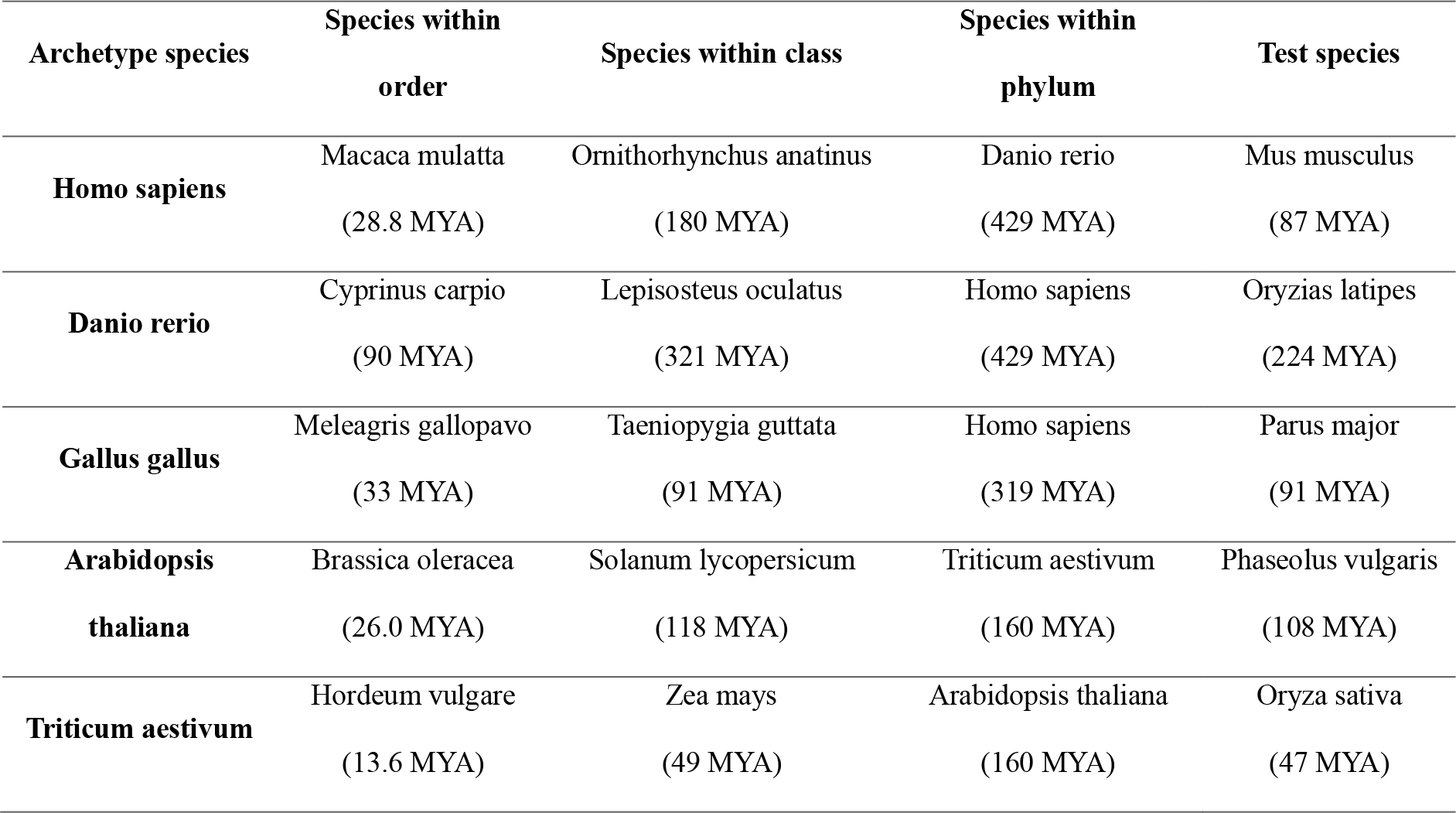
Species used for training the result filtering model and for testing performance. The numbers inside the parentheses indicate the evolutionary distances between each species and the corresponding archetype species.

**Figure 2.**
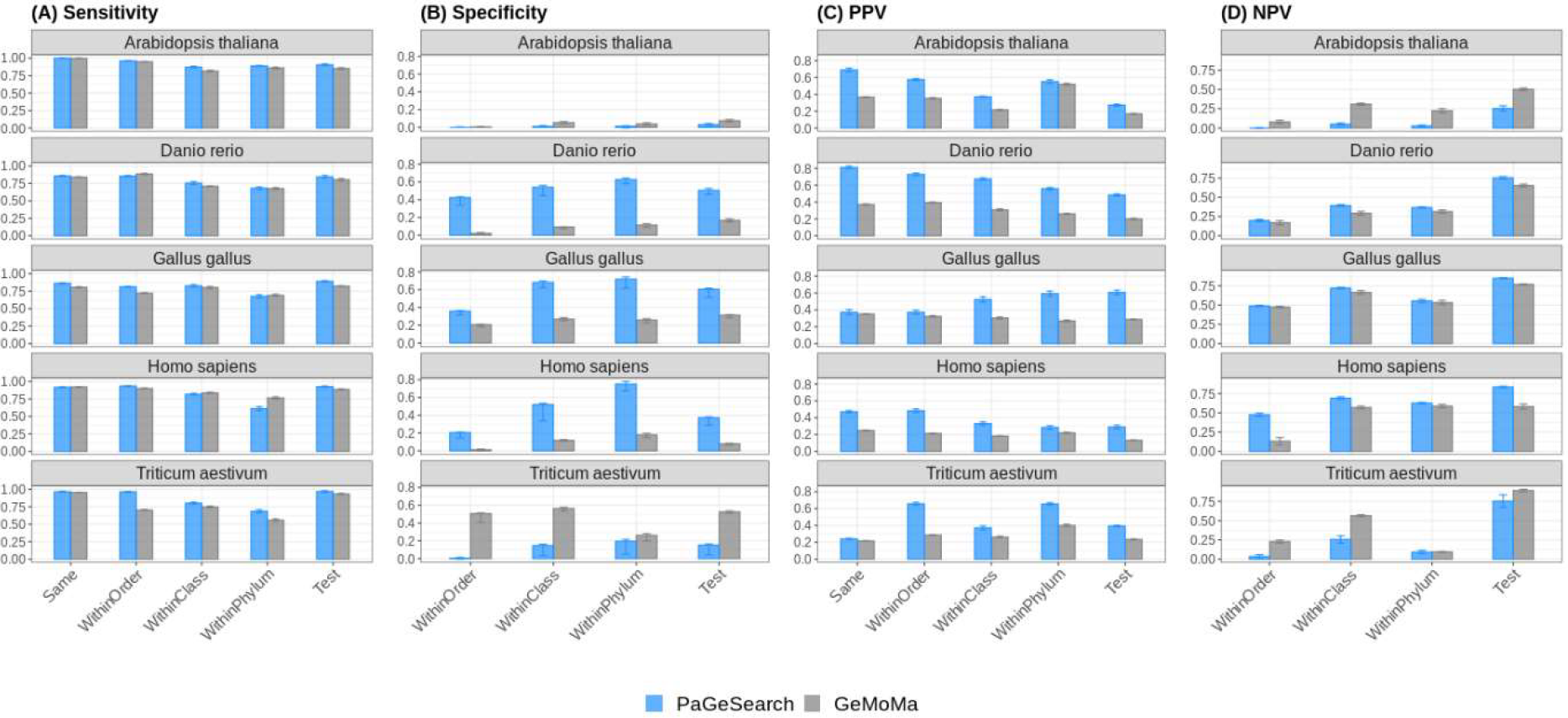
The (A) sensitivity, (B) specificity, (C) PPV, and (D) NPV of PaGeSearch and GeMoMa tested on randomly selected genes. Sensitivity and PPV evaluations were conducted on both the four species utilized for model construction and a different test species. Specificity and NPV assessments were carried out on the three species employed in model construction (excluding the same species) and a different test species.

Here, sensitivity refers to the proportion of query genes present in the genome that are successfully identified by PaGeSearch or GeMoMa in the results. On the other hand, specificity represents the proportion of query genes not present in the genome that are correctly not found by PaGeSearch or GeMoMa. PPV is the proportion of correctly identified genes among those found in the results by PaGeSearch or GeMoMa, while NPV is the proportion of genes not found in the results by PaGeSearch or GeMoMa that are actually not present in the genome.

In terms of sensitivity, PaGeSearch outperformed GeMoMa when searching for genes in the ‘Test’ species and the ‘Within Class; species. For gene searches within the ‘Same’ species, PaGeSearch demonstrated higher sensitivity, except for Homo sapiens, however the difference was marginal, less than 0.01. When tested on ‘Within Order’ species, PaGeSearch exhibited higher sensitivity, except for Danio rerio. The average sensitivity of all archetype species was 0.92 for in the ‘Test’ species, indicating its capability to identify over 90% of the genes within a given pathway or list.

The specificity of PaGeSearch consistently exceeded that of GeMoMa across all cases for Homo sapiens, Danio rerio, and Gallus gallus. In contrast, GeMoMa exhibited higher specificity for Arabidopsis thaliana and Triticum aestivum. A lower specificity suggests a higher proportion of genes that are absent from the actual genome that are erroneously listed as present in the prediction results.

PaGeSearch consistently yielded higher PPV compared to GeMoMa. This implies that PaGeSearch generates fewer false positive predictions, which can be attributed to its ability to identify the most probable ortholog for each query gene, reducing the likelihood of false positive identifications.

Like specificity, NPV was higher in animal species for PaGeSearch and higher in plant species for GeMoMa. In animals, PaGeSearch had a higher proportion of genes predicted to be negative that were actually not present in the genome, whereas in plants, this proportion was lower.

We also evaluated the extent to which predicted genes covered regions of the actual genes (Figure 3). The gene coverage was calculated as the proportion of the actual gene that intersects with the predicted gene and was obtained from only true positive predictions. The overall mean of gene coverage was 0.74 for PaGeSearch and 0.72 for GeMoMa. In more detail, the mean gene coverage for PaGeSearch when tested on the same species was 0.76, while for GeMoMa it was 0.71. When tested on different test species, the mean coverage amounted to 0.75 and 0.74 for PaGeSearch and GeMoMa, respectively. These results indicate that the majority of the gene regions are covered by the predicted regions, and PaGeSearch generally provided slightly higher coverage of the actual gene regions compared to GeMoMa.

**Figure 3.**
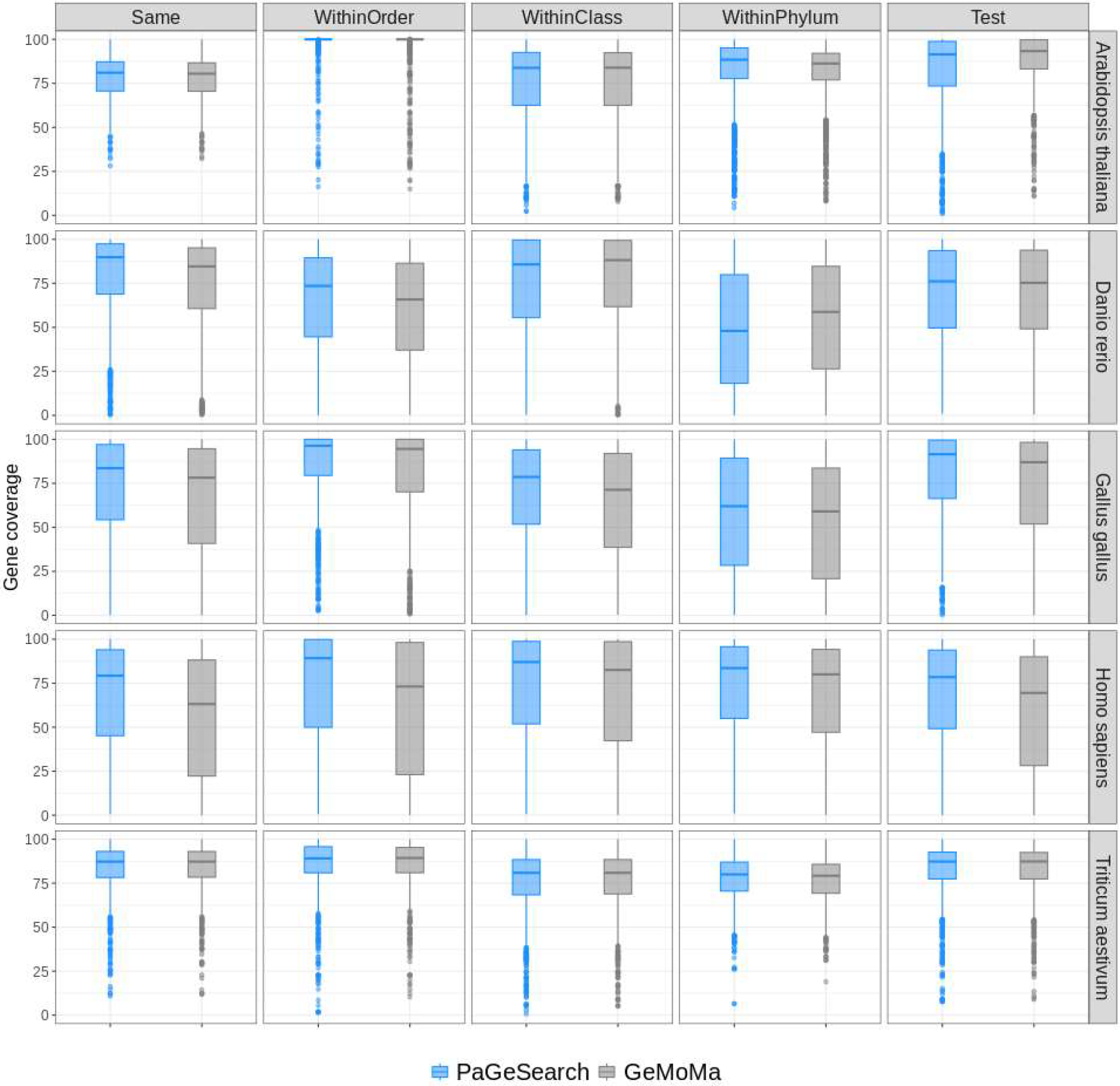
The distributions of gene coverages for true positive regions, as identified by PaGeSearch and GeMoMa, categorized by archetype species and the types of species used for validation.

Additionally, we annotated the regions identified by PaGeSearch and GeMoMa (Figure 4). The annotations encompass various gene types: “Correct genes” represent regions that coincide with the query gene regions; “Paralogous genes” correspond to regions that intersect with paralogs of the query genes; “Other genes” denote regions overlapping with protein-coding genes that are not included in the query, nor are they paralogous to it; “Pseudogenes” signify regions that overlap with pseudogene annotations; and finally, “Intergenic” refers to regions that do not exhibit overlaps with any gene annotations. Most of the false positive predicted genes were either paralogous genes or other protein coding genes in both PaGeSearch and GeMoMa.

**Figure 4.**
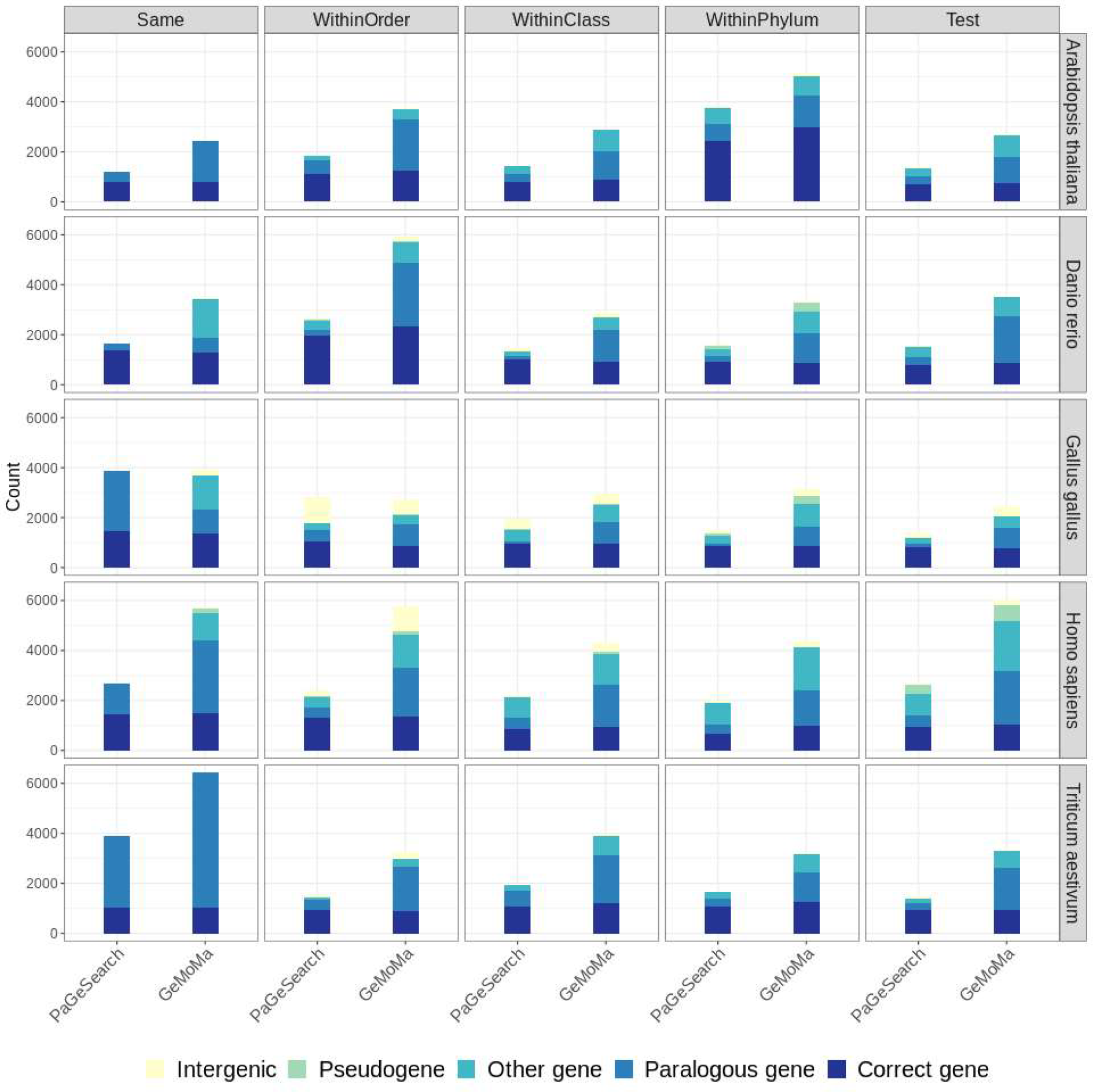
The annotations of regions identified in PaGeSearch and GeMoMa categorized by archetype species and type of species used for validation. ‘Correct genes’ are regions that overlap with the regions of the query genes; ‘paralogous genes’ are regions that overlap with paralogs of the query genes; ‘other genes’ are regions that overlap with protein coding genes that are neither correct genes nor paralogs of the query genes; ‘pseudogenes’ are regions that overlap with pseudogenes; and ‘intergenic’ are regions that do not overlap with any genes.

Overall, in plants, paralogous genes contributed significantly to false positives, while in animals, both paralogous genes and other genes were the main contributors. When PaGeSearch was tested in Arabidopsis thaliana and Triticum aestivum, 57% and 88% of the false positives were paralogous genes. On the other hand, paralogous genes compromised 42∼51% of the false positives in Danio rerio, Gallus gallus, and Homo sapiens. Lower annotation quality seems to result in a higher presence of pseudogenes and intergenic regions, particularly in bird species.

### Application of PaGeSearch to Real Pathways

We selected five common pathways that are shared between animals and plants and acquired the orthologs of genes included in these pathways using the methodology outlined in Section 2.1. These orthologs were then employed as queries. The search for orthologs was conducted within the largest taxon encompassing the archetype species, excluding the distinct test species. PaGeSearch’s performance with actual pathways was assessed in a manner analogous to the benchmark evaluation (Figure 5).

**Figure 5.**
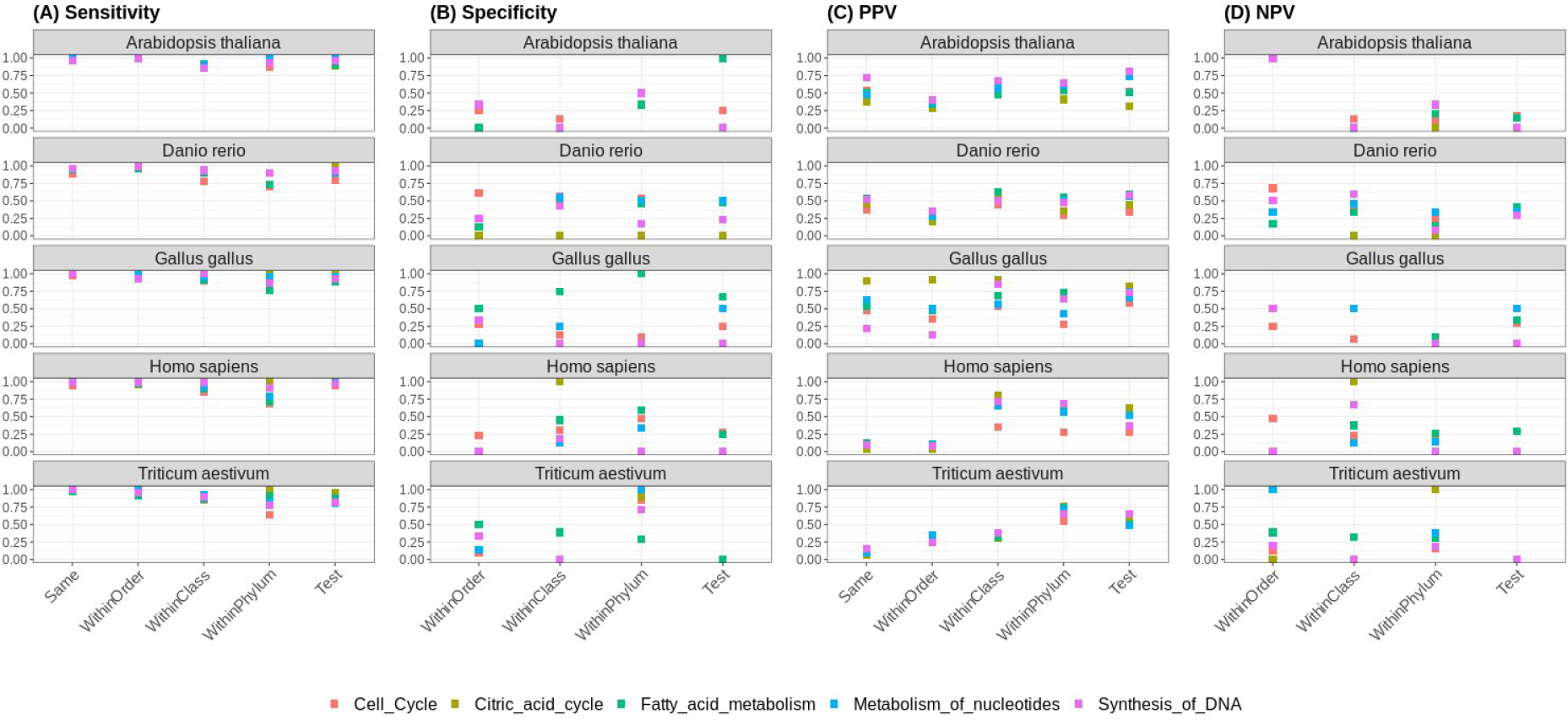
The (A) sensitivity, (B) specificity, (C) PPV, and (D) NPV of PaGeSearch tested on five common pathways. Sensitivity and PPV evaluations were conducted on both the four species utilized for model construction and a different test species. Specificity and NPV assessments were carried out on the three species employed in model construction (excluding the same species) and a different test species.

The average sensitivity across the five pathways ranged from 0.94 to 0.98 when tested on the ‘Same’ species, from 0.95 to 1 when tested on ‘Within Order’ species, from 0.91 to 0.96 when tested on ‘Within Class’ species, and from 0.90 to 0.98 when tested on ‘Test’ species. In the majority of cases, sensitivity surpassed 0.9. For the ‘Same’ species, only one pathway in Danio rerio exhibited sensitivity below 0.9, and among the ‘Test’ species, 21 out of the 25 pathways among the five archetype species displayed sensitivities above 0.9.

There were cases where specificity was unattainable because none of the genes in the pathway did not have orthologs in the genome being tested. Specificity ranged from 0 to 1, and the average specificity for the ‘Within Order’ species was 0.12, for the ‘Within Class’ species it was 0.16, and for the ‘Test’ species it was 0.22.

The PPV values ranged from 0.27 to 1 for all cases tested. The average for Arabidopsis thaliana, Danio rerio, Gallus gallus, and Homo sapiens was 0.80, 0.75, 0.89, and 0.74 respectively. For Triticum aestivum the average PPV in the ‘Same’ species was 0.29 and in other species were 0.94.

The NPV values showed a wider range when tested on the actual pathways than on random gene sets. The standard variations of NVP for common pathways tested in ‘Within Order’, ‘Within Class’, and ‘Test’ species were 0.35, 0.27, and 0.21 respectively. The average in ‘Within Order’, ‘Within Class’, and ‘Test’ species were 0.29, 0.22, and 0.17 respectively, which was lower than the NPV for random gene sets.

### Runtime

We measured the runtime of PaGeSearch for different pathways and genomes using four CPUs (Figure 6). The runtime increases as the genome size and the number of genes in the pathway increases. For the smallest genome used for validation, Arabidopsis thaliana (135Mb), it took 33 to 164 seconds to identify 11 to 67 genes in the pathway. For the largest genome, Triticum aestivum (17Gb), the runtime ranged from 2923 to 3756 seconds, in order to find 31 to 187 genes in the pathway.

**Figure 6.**
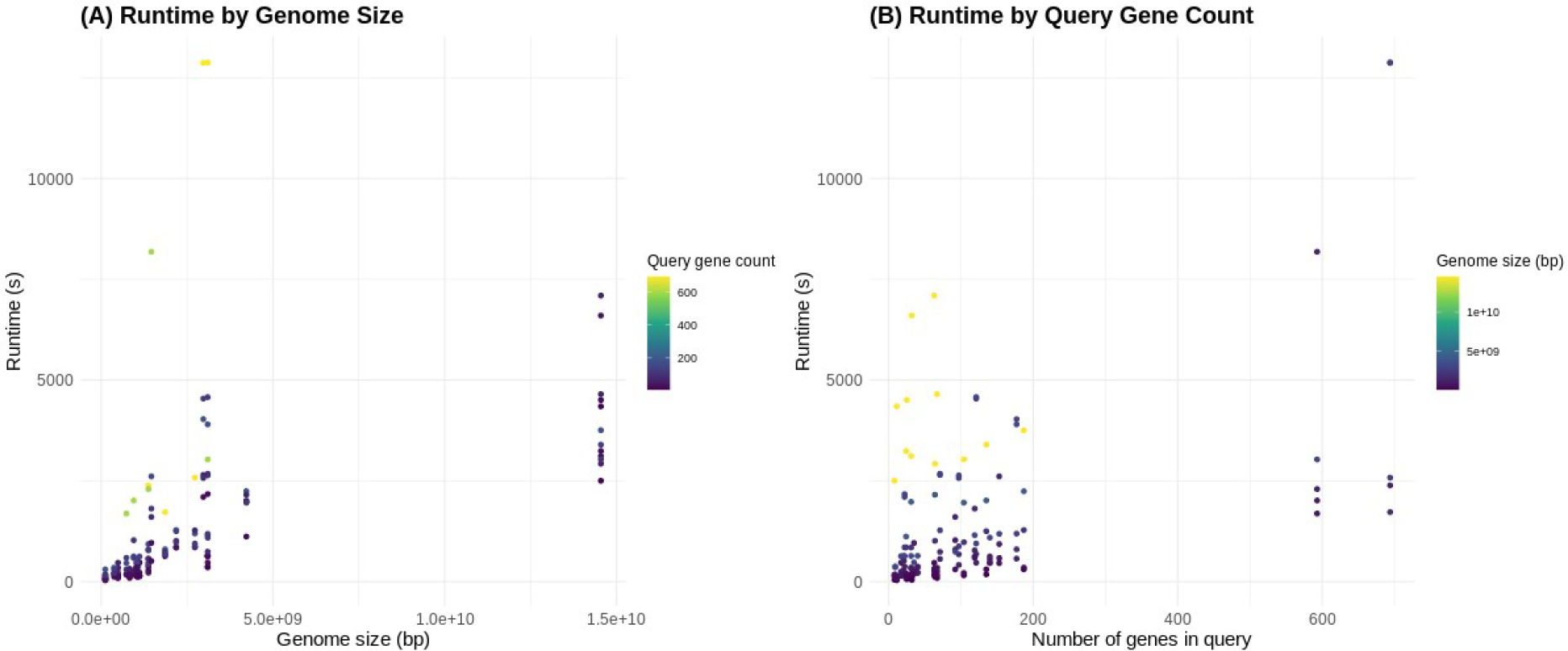
Runtime of PaGeSearch by Genome size and query gene count. (A) Runtime increases as the size of the genome being searched increases. The unit for runtime is seconds, and the unit for genome size is base pairs (bp). (B) Runtime increases as the query gene count increases. The unit for runtime is seconds, and the query gene count is measured in the number of genes.

## DISCUSSION

PaGeSearch uses a multi-step approach, first employing similarity searches to pinpoint candidate regions, and then carrying out gene prediction within these relatively small genomic fractions. This makes it more efficient than methods that annotate entire genomes. For instance, BRAKER2 processed a 1 Mb nucleotide segment of Arabidopsis thaliana’s genome in 20 to 35 minutes on 4 CPUs, while PaGeSearch can find 67 genes in the Arabidopsis thaliana’s genome in 3 minutes on 4 CPUs (Brůna et al. 2021).

The final step of PaGeSearch is filtering hits based on similarity search scores, gene prediction statistics, and alignment scores of the predicted proteins and query proteins. PaGeSearch aims to find the most likely orthologous genes of the genes in the query. This approach enables PaGeSearch to maintain relatively higher PPV than other tools. Regarding PPV, PaGeSearch consistently outperforms GeMoMa.

In general, as the evolutionary distance increases, sensitivity tends to decrease whereas specificity tends to increase in both PaGeSearch and GeMoMa. This trend is consistent across the hierarchical classifications of ‘Same’ species, ‘Within Order’ species, ‘Within Class’ species, and ‘Within Phylum’ species. ‘Test’ species yields sensitivity and specificity levels intermediate to those of ‘Within Order’ species and ‘Within Class’ species. However, deviations from these patterns arise, potentially influenced by discrepancies in genome annotation quality. Well-annotated genomes may manifest higher sensitivity and specificity, while genomes with less robust annotations could result in false positives due to unannotated genes or false negatives owing to incompletely annotated genes. NPV tends to increase with the evolutionary distance, although this is not always the case. Both sensitivity and NPV are likely influenced not only by the quality of annotation but also by the quality of ortholog definitions; if orthologs between the archetype species and the tested species are defined properly, more false positives and false negatives may result.

Also, when dealing with query genes that have a large number of paralogous genes within the genome, the specificity, PPV and NPV of PaGeSearch can be compromised. The low specificity and NPV of PaGeSearch in Arabidopsis thaliana and Trivicum aestivum can be attributed to the detection of many paralogs, as most of the false positives were paralogous genes (Figure 4). This is relevant since flowering plants like these typically have a higher number of paralogs due to frequent and recent whole-genome duplication events. (Soltis et al. 2015). PaGeSearch operates in two main steps: it first uses a sequence similarity search to identify all possible regions, and then filters these based on neural network models. While this approach allows for high sensitivity, it may also result in identifying some false positives, such as paralogs, whose sequences are similar to those of the query genes.

To elaborate, when using Arabidopsis thaliana as the archetype, its compact genome and well-annotated status, where paralogs and genes with the same function are specifically annotated to have separate names, could lead to fewer true negatives being identified. This affects the performance metrics of both PaGeSearch and GeMoMa manifesting as lower specificity for each tool when working with this species. In contrast, Triticum aestivum has a hexaploid genome, inherently containing more copies of the same genes. PaGeSearch relies on sequence similarity for its search criteria, while GeMoMa uses reference sequence and transcript annotation information. As a result, PaGeSearch is more susceptible to reduced true negative identification due to polyploidy, while GeMoMa is less affected, leading to higher sensitivity in Triticum aestivum.

The PPV patterns of PaGeSearch do not correlate with evolutionary distances, suggesting that other factors are at play. High PPV is more likely when the genome has few regions similar to the query and when there are minimal ties in prediction probability. These aspects may contribute to PPV patterns that do not align straightforwardly with evolutionary distances.

When validated against actual pathways, the mean sensitivity for Arabidopsis thaliana, Danio rerio, Gallus gallus, Homo sapiens, and Triticum aestivum was 0.96, 0.91, 0.95, 0.94, and 0.94 respectively. This indicates that PaGeSearch can successfully identify more than 90% of genes within real pathways, on average. The overall average specificity across species ranged from 0 to 1, and the average PPV ranged from 0.29 to 0.95. When applied to common pathways, the sensitivity was higher or comparable to that of the random test set, while the specificity and PPV were somewhat lower. Real pathways may contain similar genes within the query. Furthermore, since we tested essential pathways, there may be a number of genes within the pathway query that share high similarity with other genes in the genome. These may have led to the lower specificity, PPV, and NPV in pathway results compared to random test set validations, as the existence of genes that have similar sequences with the query genes both within the query and in the genome may yield more false positives. Although the performance demonstrates increased variability when assessed within actual biological pathways, PaGeSearch maintains high sensitivity and the specificity, PPV, and NPV fall within a range that makes the tool suitable for practical applications in most cases.

PaGeSearch can accurately identify genes within pathways and demonstrate its utility across various species within the same classes as the five archetype species. Currently, PaGeSearch can be applied to a class comprising five archetype species. However, we have plans to extend its applicability to a broader range of species, such as insects. This update aims to provide a more comprehensive coverage across diverse taxa. The versatile pipeline offered by PaGeSearch will enable similar analyses to be conducted on species beyond this scope, ensuring its utility for a wide variety of organisms.

In conclusion, PaGeSearch offers a convenient pipeline that can download the sequences of the genes in a pathway and search them in the genome, when only the Pathway ID and genome assembly are provided. PaGeSearch reduces the effort required of searching for gene lists and their corresponding sequences within a pathway, enabling relatively straightforward analysis. Furthermore, the high sensitivity ensures that the majority of genes within the pathway can be identified, while PaGeSearch maintains a shorter runtime compared to other gene prediction tools. Rather than focusing solely on homologs, PaGeSearch seeks the most probable orthologs, resulting in lower false positive rates. Because it identifies gene-predicted regions exclusively, its interpretation is simpler compared to similarity search tools such as BLAST. PaGeSearch offers a convenient means for comparative analyses of specific pathways across multiple species, especially when the genome annotation is unavailable or the quality of annotation varies among species.

## METHODS

PaGeSearch integrates multiple tools to search regions that are both similar to given gene sequences and are predicted to be genic (Figure 1). First, seed regions are discovered through a sequence similarity search, followed by the grouping and extension of these regions. Within these candidate regions, ab initio gene prediction is employed to detect genes. Finally, the predicted gene sequence and query gene sequence are aligned, and the results are filtered according to the scores calculated by a neural network model evaluating the similarity search scores and the protein alignment scores.

The models used for result filtering was constructed based on five archetype species and are designed to be generally applicable to other species within the same taxonomic classes: Homo sapiens (mammals), Danio rerio (fish), Gallus gallus (birds), Arabidopsis thaliana (eudicotyledons), and Triticum aestivum (liliopsida). Similarly, other parameters within the pipeline have been adjusted to be applicable to species that fall within the same classes as these five archetype species. The pipeline was implemented using Python programming language and default parameters were applied for all software components unless otherwise specified.

PaGeSearch also provides a data collection pipeline based on the Reactome pathway database (Gillespie et al. 2022). The pipeline downloads protein sequences of all genes in a given pathway name, as well as the ortholog sequences of those genes for a specific taxon. The sequences are sourced from the Ensembl database (Cunningham et al. 2022) and can be used directly as input for the gene searching pipeline.

### Data Collection

As an initial step for pathway gene searching, users can obtain the sequences of genes associated with their pathway of interest. Users have the option to provide either a list of specific genes or Reactome pathway names. When given the pathway name and species names, the set of Ensembl gene IDs related to the pathway is sourced from the Reactome database (Gillespie et al. 2022). Subsequently, gene names and their orthologs are gathered from the Ensembl REST API at rest.ensembl.org (Ensembl Release 109, Feb 2023) (Yates et al. 2015). equences of orthologs, including their isoforms, are accessed via the Ensembl REST API and are stored in individual FASTA files.

### Candidate Region Identification Through Sequence Similarity Search

First, PaGeSearch consolidates the FASTA files of all genes from the provided list into a unified FASTA file and conducts quality control, purging any sequences with ambiguous amino acids (BJOUZ). To boost computational efficiency, only distinct sequences are retained, achieved using SeqKit rmdup (Shen et al. 2016). We utilize MMseqs2 for the sequence similarity search (Steinegger and Söding 2017). This involves creating a protein database with the query genes and a nucleotide database of the target genome, followed by executing a tblastn search.

### Candidate Region Grouping and Extension

MMseqs2 outputs are grouped and expanded for subsequent gene prediction. In particular, High-scoring Segment Pairs (HSPs) are categorized by gene. HSP regions within a singular gene and strand, which are separated by less than 100,000 bp, are amalgamated into one extensive region. Each of these regions is then augmented by a margin of 10,000 bp on either side. These enlarged regions are earmarked as potential regions for the ensuing analyses. We generated a summary of sequence similarity search metrics from these candidate regions for future result filtering. Aligned query gene sequences are organized by identical positions and score, retaining only the top five sequences for matches at identical locations for continued analysis. To acquire these sequences, we utilized SAMtools faidx (Li et al. 2009) and BEDTools getfasta (Quinlan and Hall 2010).

### Making Protein Hints

For improving gene prediction accuracy, PaGeSearch deploys exonerate (with the protein2genome model) and an internal perl script from exonerate, exonerate2hints.pl, to make protein hints (Slater and Birney 2005). The source data consists of sequences from the query genes and the candidate regions described in the previous section.

### Ab initio Gene Prediction

Ab initio gene prediction on the candidate regions are performed by AUGUSTUS (Stanke et al. 2006). PaGeSearch searches for complete gene regions, incorporating the protein hints file created in the previous section for the corresponding species model. The protein sequences of the predicted gene regions are extracted using getAnnoFasta.pl.

### Protein Sequence Alignment

PaGeSearch aligns the protein sequences of the query gene sequences to the predicted gene sequences, to validate the gene prediction findings. The ungapped model from Exonerate was utilized to find the best alignment for each predicted gene sequence and obtain the alignment statistics (Slater and Birney 2005).

### Result Filtering

PaGeSearch filters the gene prediction outputs by employing a pre-trained model that integrates metrics from sequence similarity search, gene prediction, and protein alignment. We produced training datasets for each species using PaGeSearch, bypassing the result filtering phase on a selection of thoroughly annotated genomes. We derived all peptide sequences of the archetype species, excluding sequences present in the shortest and longest 10% to omit outliers. Next, we collected genome sequences from four species for each archtype species with varying evolutionary distances, so that we can obtain a comprehensive training dataset enabling the general application of the constructed model. The four species were as follows; 1) the archetype species, 2) close species within the same order, 3) species from the same class but a different order, and 4) distant species within the same phylum (Table 1, Supplementary Table 1). The evolutionary distances showcased in Table 1 were extrapolated from the TimeTree database (Hedges et al. 2015).

We labeled the outcomes of gene searching as 1 if there was an overlap with the query and 0 otherwise. To balance the data, synthetic minority over-sampling technique (SMOTE) was performed so that the ratio of label 1 and 0 would be 1:1 using python library imbalanced-learn (Lemaître et al. 2017). A neural network model was trained using the features in Supplementary Table 2 to and the labels of the outcomes. The neural network model was constructed using the Keras Sequential API (Chollet and et al. 2015). The model architecture comprised three layers. The initial layer is a dense layer with 4 neurons, followed by another dense layer with 2 neurons. The final layer is a dense layer with 1 neuron, applying a sigmoid activation function, designed for binary classification tasks. To optimize the model, the Adam optimizer was employed, with a learning rate of 10^−5^, and the model was compiled with binary cross-entropy as the loss function.

The predictions were sorted based on the probability of being labeled as 1. When multiple genes were predicted in the same genomic region, only the one with the highest probability was retained. Subsequently, if the same gene was predicted in multiple genomic regions, the prediction with the highest probability was selected. In cases where probabilities were equal or very similar (with a difference of 0.05 or less), multiple predictions were considered. Afterwards, only predictions with probabilities greater than or equal to a threshold were selected for further analysis. This approach ensures that the most confident and reliable predictions are considered while accounting for cases where probabilities are close or equal.

### Validation

To assess the performance of PaGeSearch, we conducted evaluations in two scenarios: 1) searching for pathway genes within the genomes of the four species used to construct the models, and 2) searching for pathway genes within the genome of a different test species (Table 1). To evaluate both sensitivity and specificity, we randomly selected 200 genes from the archetype species. Among these, 100 genes had orthologs in the test species, while the remaining 100 genes did not have orthologs. If there was overlap between the regions identified by PaGeSearch and the actual query gene regions, these were considered true positives. Regions identified by PaGeSearch that did not overlap with the query were considered false positives. Query regions not present in the PaGeSearch results were classified as false negatives. We considered query genes that were not present in either the searched genome or the PaGeSearch results to be true negatives. The presence of a query gene in a different genome being searched was determined by the existence of orthologous genes. This process was repeated five times, and the definition of orthologs was based on the Ensembl database.

## SOFTWARE AVAILABILITY

PaGeSearch source code is available at https://github.com/Sohyoung/PaGeSearch.

## ACKNOWLEDGMENTS

We would like to express our gratitude to eGenome Inc. for their support throughout the course of this research.

